# SARS-CoV-2 Omicron BA.1 and BA.2 are attenuated in rhesus macaques as compared to Delta

**DOI:** 10.1101/2022.08.01.502390

**Authors:** Neeltje van Doremalen, Manmeet Singh, Taylor A. Saturday, Claude Kwe Yinda, Lizzette Perez-Perez, W. Forrest Bohler, Zachary A. Weishampel, Matthew Lewis, Jonathan E. Schulz, Brandi N. Williamson, Kimberly Meade-White, Shane Gallogly, Atsushi Okumura, Friederike Feldmann, Jamie Lovaglio, Patrick W. Hanley, Carl Shaia, Heinz Feldmann, Emmie de Wit, Vincent J. Munster, Kyle Rosenke

## Abstract

Since the emergence of SARS-CoV-2, five different variants of concern (VOCs) have been identified: Alpha, Beta, Gamma, Delta, and Omicron. Due to confounding factors in the human population, such as pre-existing immunity, comparing severity of disease caused by different VOCs is challenging. Here, we investigate disease progression in the rhesus macaque model upon inoculation with the Delta, Omicron BA.1, and Omicron BA.2 VOCs. Disease severity in rhesus macaques inoculated with Omicron BA.1 or BA.2 was lower than those inoculated with Delta and resulted in significantly lower viral loads in nasal swabs, bronchial cytology brush samples, and lung tissue in rhesus macaques. Cytokines and chemokines were upregulated in nasosorption samples of Delta animals compared to Omicron BA.1 and BA.2 animals. Overall, these data suggests that in rhesus macaques, Omicron replicates to lower levels than the Delta VOC, resulting in reduced clinical disease.

---

SARS-CoV-2 is under constant evolutionary pressure. The unprecedented speed and volume of whole-genome sequencing employed during the pandemic has allowed for near real-time surveillance of amino acid substitutions. The close surveillance of virus genomes for such substitutions additionally led to early detection and analysis of variants of concern (VOCs) (1). A variant is deemed a VOC when it displays evidence for increased transmissibility, increased disease severity, or decreased effectiveness of available diagnostics, vaccines, and therapeutics (2). The first recognized VOC was detected in September 2020 (3) and was designated Alpha. Thus far, five VOCs have been identified: Alpha (Pango lineage B.1.1.7), Beta (B.1.351), Gamma (P.1), Delta (B.1.617.2), and Omicron (B.1.1.529, which includes BA.1, BA.2, BA.3, BA.4, BA.5, and all its descendent lineages). The Delta VOC was first detected in the spring of 2021 in India. It spread very quickly on a global level, replacing the Alpha variant in the United Kingdom and United States (3–5). Delta is characterized by a number of key substitutions, such as the L452R and P681R substitutions in the S protein (6). The Omicron VOC was then detected in November 2021 in South Africa, and subsequently replaced the Delta VOC. Omicron is characterized by >30 substitutions in the S protein (6).

Studies aiming to identify the evolutionary advantages of each VOC in the human population are complex, due to population-wide confounding factors such as previous SARS-CoV-2 infections and vaccine coverage. Animal models allow us to study pathogenesis and compare viral replication kinetic in naïve animals, thereby circumventing these confounders. We previously utilized the rhesus macaque model to examine differences in pathogenicity between an ancestral strain (Wuhan-like) with the D614G mutation, the Alpha VOC, and the Beta VOC and showed that inoculation with the Beta VOC resulted in lower clinical scores, lower lung virus titers, less severe lung lesions, and lower cytokine and chemokines in the bronchoalveolar lavage (7). In the current study, we aim to extend this data set to include the Delta, Omicron BA.1, and Omicron BA.2 VOCs in naïve rhesus macaques.

## Results

In this study, we compared three different SARS-CoV-2 isolates: the Delta AY.106 VOC (hCoV-19/USA/MD-HP05647/2021, EPI_ISL_2331496); the Omicron BA.1 VOC (hCoV- 19/USA/GA-EHC-2811C/2021, EPI_ISL_7171744), and the Omicron BA.2 VOC (hCoV-19/Japan/UT-NCD1288-2N/2022, EPI_ISL_9595604). All stocks were sequenced and no substitutions in the S protein, as compared to published sequences, were found. For a comparison of S protein sequences, refer to **Table S1**.

To determine the entry profile of the respective VOCs, we compared the entry of pseudotyped vesicular stomatitis virus (VSV) particles expressing the S protein of Wuhan1 virus to particles expressing the S protein of the Delta, Omicron BA.1, and Omicron BA.2 VOC into baby hamster kidney cells (BHKs) expressing either the human or rhesus ACE2. Entry was observed under all conditions but was significantly less efficient for the Omicron VOCs compared to the Delta VOC, both with human and rhesus ACE2 (**Figure S1**).

### Reduced clinical signs in Omicron-inoculated rhesus macaques

Three groups of six rhesus macaques were inoculated intranasally and intratracheally with a total dose of 2 × 10^6^ median tissue culture infective dose (TCID50) of one of the SARS-CoV-2 VOCs. Although animals in all three groups showed mild signs of disease after challenge, inoculation with Delta resulted in noticeable higher clinical scores than inoculation with Omicron BA.1 and BA.2, a result that was only statistically significant for Omicron BA.1 (**Figure 1A**). Most animals in all three groups had days with reduced appetite throughout the study. However, respiratory signs were significantly different between groups: they were observed in four animals inoculated with Delta, only one animal inoculated with Omicron BA.2, and no animals inoculated with Omicron BA.1 (**Figure 1 B-C**). Radiographs collected on all exam days were analyzed for the presence of pulmonary infiltrates. Most animals in all three groups did not present with pulmonary infiltrates, except for one animal each in the Delta and Omicron BA.1 challenged groups. (**Figure S2A**). No major changes were observed in the body weight or temperature of NHPs during the study (**Figure S2 B-C**).

**Figure 1.**
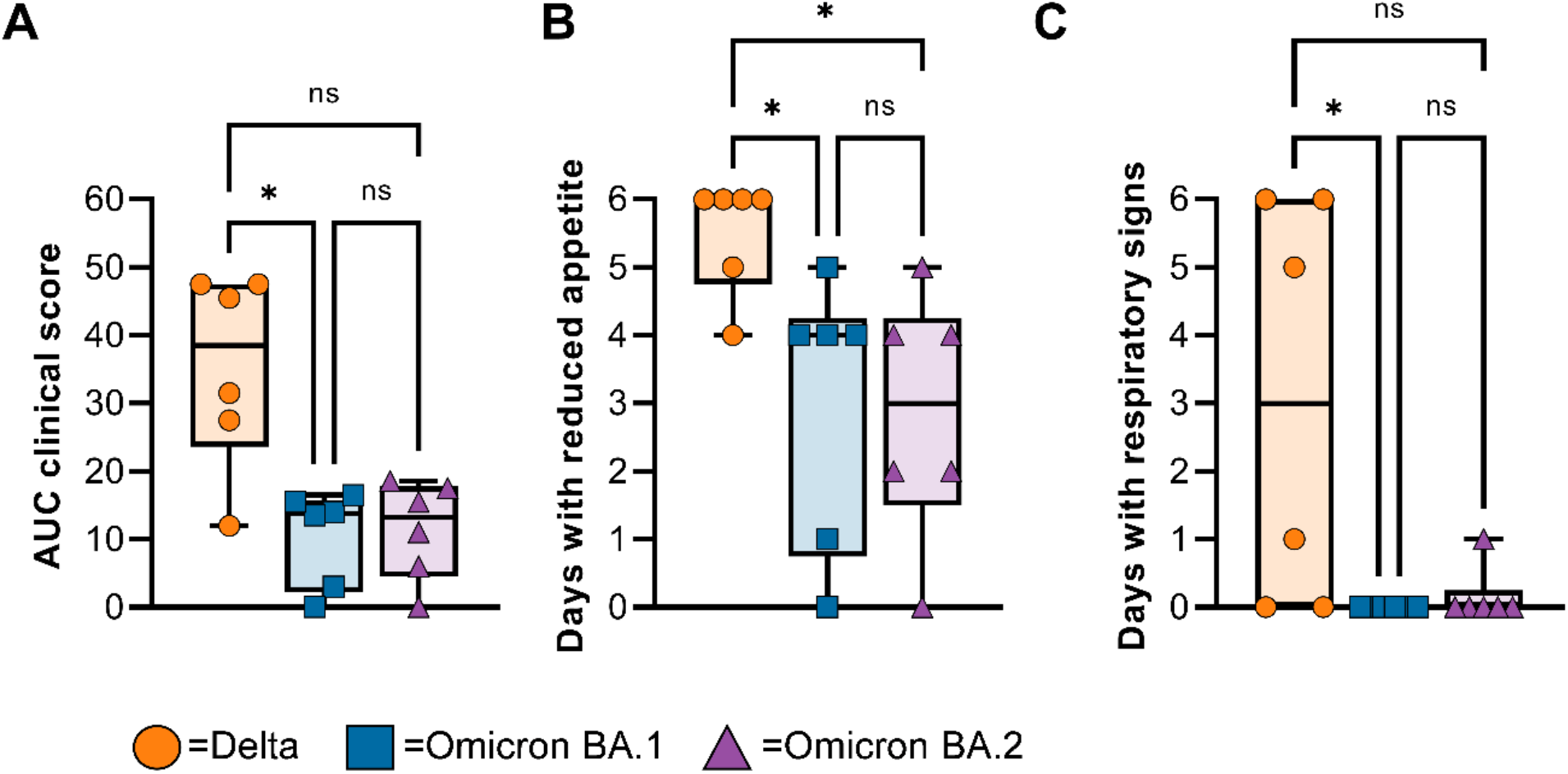
Rhesus macaques inoculated with Omicron display milder disease than animals inoculated with Delta. Three groups of six adult rhesus macaques were challenged with SARS-CoV-2 VOCs Delta (orange circles), Omicron BA.1 (blue squares), or Omicron BA.2 (purple triangles). (A) Daily scores of disease signs for each animal were utilized to calculate one area-under-the-curve number per animal and displayed in a minimum-to-maximum boxplot. (B) The days in which reduced appetite was noted are totaled per animal and shown as a boxplot (minimum-to-maximum). (B) The days in which respiratory signs were noted are totaled per animal and shown as a boxplot (minimum to maximum). (C) Minimum-to-maximum boxplot of radiographs taken on exam days. Individual lobes were scored by a clinical veterinarian according to a standard scoring system and totaled. Statistical analysis was performed using a Kruskal-Wallis test with Dunn’s multiple comparisons, * = p value <0.05.

### Reduced shedding after Omicron BA.1 or BA.2 inoculation

Nasal swabs were collected at 0-, 2-, 4-, and 6-days post inoculation (dpi) and analyzed for the presence of viral genomic RNA (gRNA) and subgenomic RNA (sgRNA). The amount of viral gRNA detected in nasal swabs from Delta animals was significantly higher than that detected in nasal swabs from Omicron BA.1 or BA.2 animals (**Figure 2A**). Similar differences were also observed in the amount of sgRNA found in nasal swabs between groups, but significance was only found 2-dpi between Delta and Omicron BA.1, and on 4- and 6-dpi between Delta and Omicron BA.2 (**Figure 2B**). For each animal, the area under the curve was calculated as a measure of the total amount of viral gRNA and sgRNA shed between 2- and 6-dpi. Animals inoculated with Delta shed significantly more gRNA than Omicron BA.1 inoculated animals, and more gRNA and sgRNA than Omicron BA.2 inoculated animals (**Figure 2A-B**). Oropharyngeal and rectal swabs were also obtained on each exam day. Presence of viral RNA in these samples was limited, compared to nasal swabs. The only significant difference between groups was found 2-dpi in gRNA in rectal swabs, where Delta animals shed significantly more than Omicron BA.1 or BA.2 animals. No viral RNA was detected in blood samples on any exam day (**Figure S3**).

**Figure 2.**
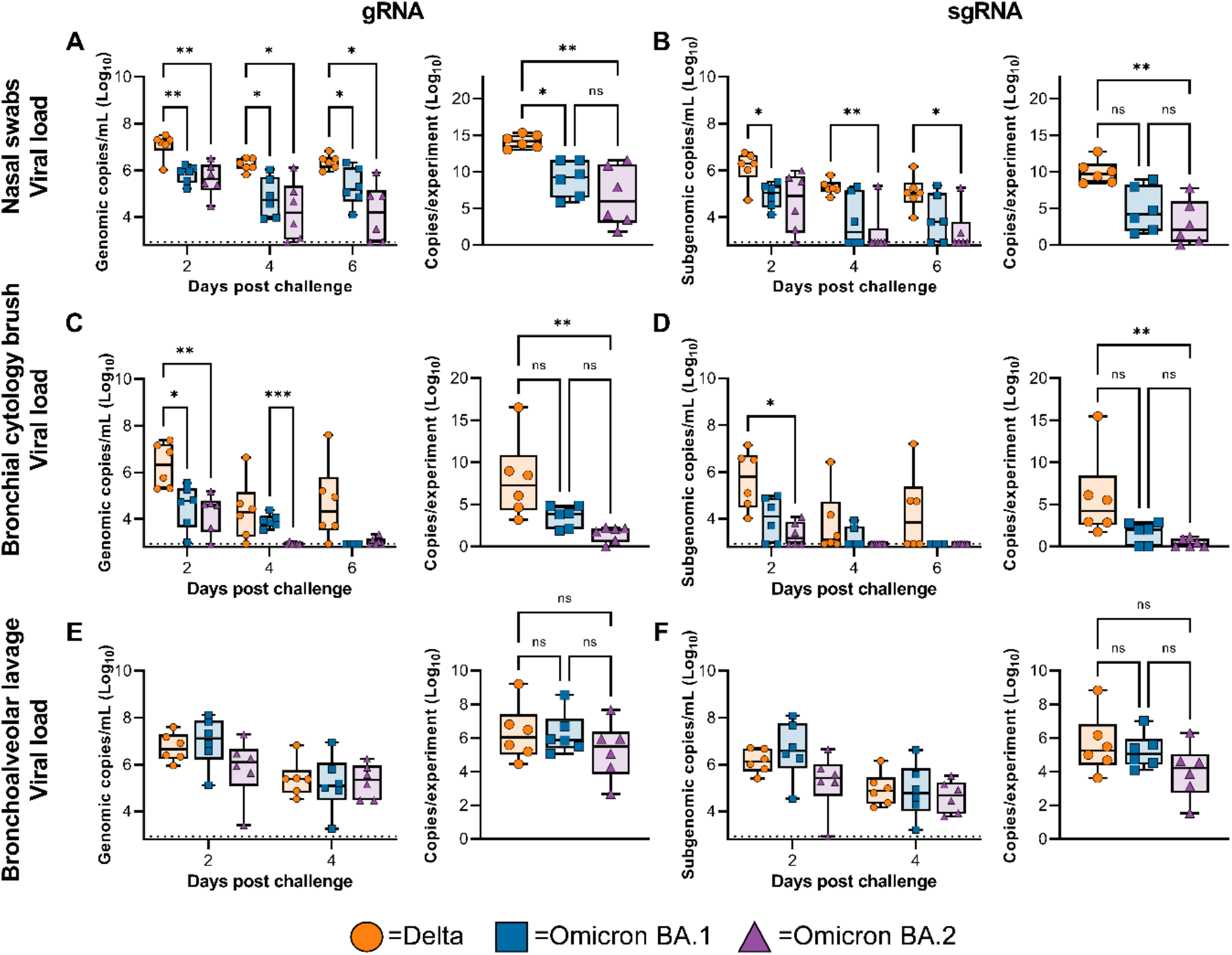
Viral load from the respiratory tract is lower in animals challenged with Omicron compared to Delta. Boxplot (minimum to maximum) of viral loads over time (left panel) and total amount of RNA detected throughout the experiment (area under the curve, right panel) in nose swabs (A-B), BCBs (C-D), and BAL fluid (E-F) taken on 2-, 4-, and 6-dpi (nose swabs and BCBs only). Statistical significance was determined via a two-way ANOVA with the Geisser-Greenhouse correction followed by the Tukey test for multiple comparisons (viral RNA per day) or via a Kruskall-Wallis test followed by Dunn’s test for multiple comparisons (area under the curve).

### Reduced virus replication in the lower respiratory tract of rhesus macaques inoculated with Omicron BA.1 and BA.2

Bronchoalveolar lavage (BAL) and bronchial cytology brush (BCB) samples were collected on 2-, 4-, and 6-dpi (BCB only) and analyzed for the presence of gRNA and sgRNA. Viral load in BAL and BCB samples were highest on 2-dpi and declined by 4- and 6-dpi (**Figure 2C-F**). As seen in the nasal swabs, less viral RNA was detection in BCB samples in animals inoculated with Omicron BA.1 or BA.2 compared to Delta (**Figure 2C-D**). In contrast, no significant differences between groups were detected in the amount of viral RNA detected in BAL samples (**Figure 2E-F**). At 6 dpi, animals were euthanized, and tissues were collected, including tissues from the upper and lower respiratory tract and intestinal tract. Although significant differences in the amount of virus detected in nasal swabs were found, the amount of viral RNA in nasal turbinates was not significantly different between groups (**Figure 3A**). In contrast, viral RNA in lung tissue was significantly lower in animals inoculated with Omicron BA.1 and BA.2 compared to Delta (**Figure 3B**). Additional tissue samples were analyzed and where positive, showed a higher gRNA and sgRNA load in Delta inoculated animals compared with Omicron BA.1 and BA.2 (**Figure S4**).

**Figure 3.**
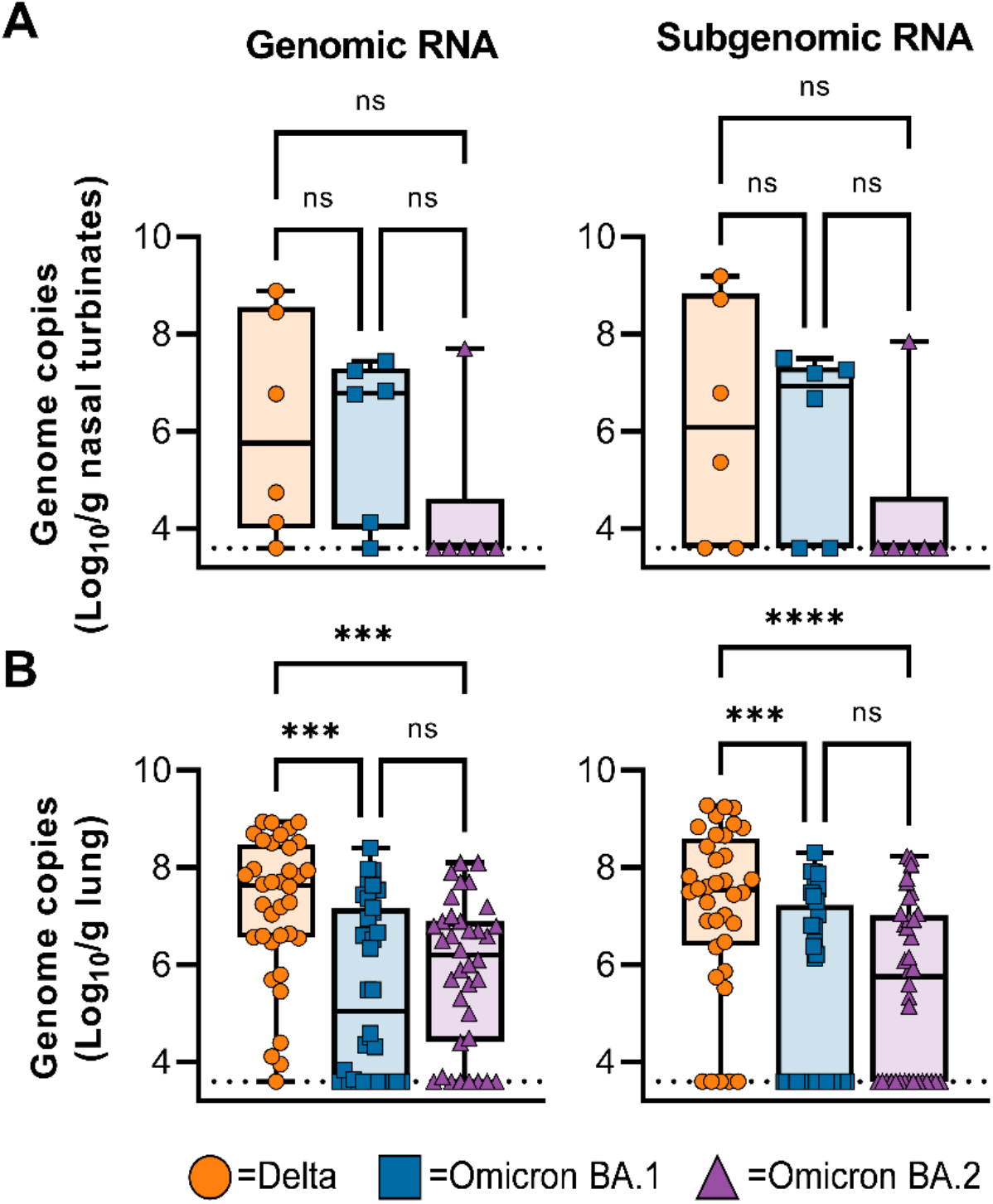
Viral loads are lower in lung tissue, but not nasal turbinates, of animals inoculated with Omicron compared to Delta on 6-dpi. (A) Boxplot (minimum to maximum) of gRNA (left panel) and sgRNA (right panel) detected in nasal turbinates. (B) Boxplot (minimum to maximum) of gRNA (left panel) and sgRNA (right panel) detected in lung tissue (shown are all six lung lobes per animal, totaling 36 samples per group). (C) Boxplot (minimum to maximum) of number of lung lobes positive for gRNA or sgRNA per animal (maximum of 6). Statistical significance was determined via a Kruskall-Wallis test followed by Dunn’s test for multiple comparisons. *** = p-value < 0.001. **** = p-value < 0.0001.

### Viral loads in respiratory tract of NHPs challenged with D614G, Alpha, and Beta variants are similar to Omicron BA.1 and BA.2, but clinical scores are higher

Compared to a previous study using the same methods and readouts (7), both Omicron BA.1 and BA.2 had lower clinical scores than D614G, Alpha, and Delta, but not Beta animals (**Figure S5A**). In nasal swabs and BCBs, viral sgRNA load was higher in Delta animals than several other variants (**Figure S5B**). In contrast, viral load in BCBs of Omicron BA.2 animals was significantly lower than those of Beta animals (**Figure S5C**). In lung tissue, samples obtained from Delta-inoculated animals were significantly higher than all other variants (**Figure S5E**). No significant differences were observed between groups in BAL samples (**Figure S5D**) or nasal turbinates (**Figure S5F**).

### Omicron BA.1 and BA.2 inoculation caused decreased respiratory pathology

In nasal turbinates, minimal-to-moderate inflammation was observed and consisted of a submucosal infiltrate of neutrophils, macrophages, and lymphocytes which infiltrated the overlaying mucosa and were interspersed with individual and small clusters of necrotic cells. SARS-CoV-2 antigen in the nasal turbinates was extremely rare and was detected in three out of six Delta challenged animals, one out of six Omicron BA.1 challenged animals, and one out of six Omicron BA.2 challenged animals within both respiratory and olfactory epithelium (**Figure 4A-B, Figure 5A-B, Figure 6A**). It is unknown as to what extent the inflammation in the turbinates may be attributable to viral challenge or is background inflammation, as SARS-CoV-2 antigen was found in both inflamed and non-inflamed tissue sections.

**Figure 4.**
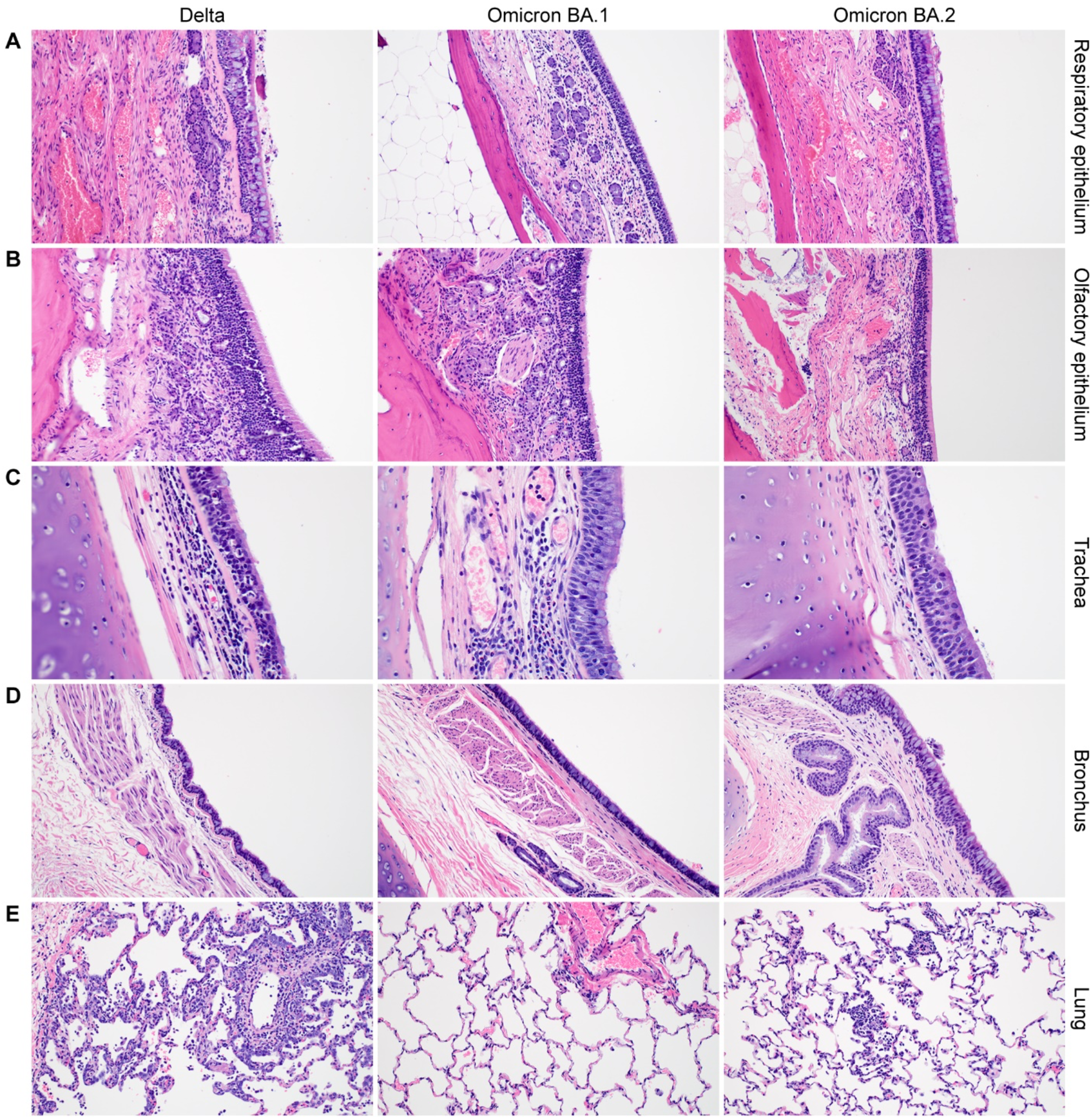
Histologic findings in the respiratory tract of SARS-CoV-2 challenged rhesus macaques at 6 dpi. (A-B). All: Minimal to mild respiratory and olfactory epithelial submucosal inflammation. (C) Delta: Tracheal inflammation is moderate and present in the submucosa and mucosa along with a few singular necrotic mucosal cells. Omicron BA.1: Inflammation is mild and limited to the submucosa. Omicron BA.2: No significant findings. (D) All: Bronchial mucosa and submucosa with no significant inflammation or necrosis. Bronchial inflammation was very rare and, when present, graded as minimal to mild. (E) Delta: Typical SARS-CoV-2 pneumonia at 6 dpi, including perivascular inflammation, thickened alveolar septa, and inflammatory cells within the alveolar lumina. Omicron BA.1: No significant findings. Omicron BA.2: Rare focus of minimal inflammation. Magnification A, B, D, E 200x; C 400x.

**Figure 5.**
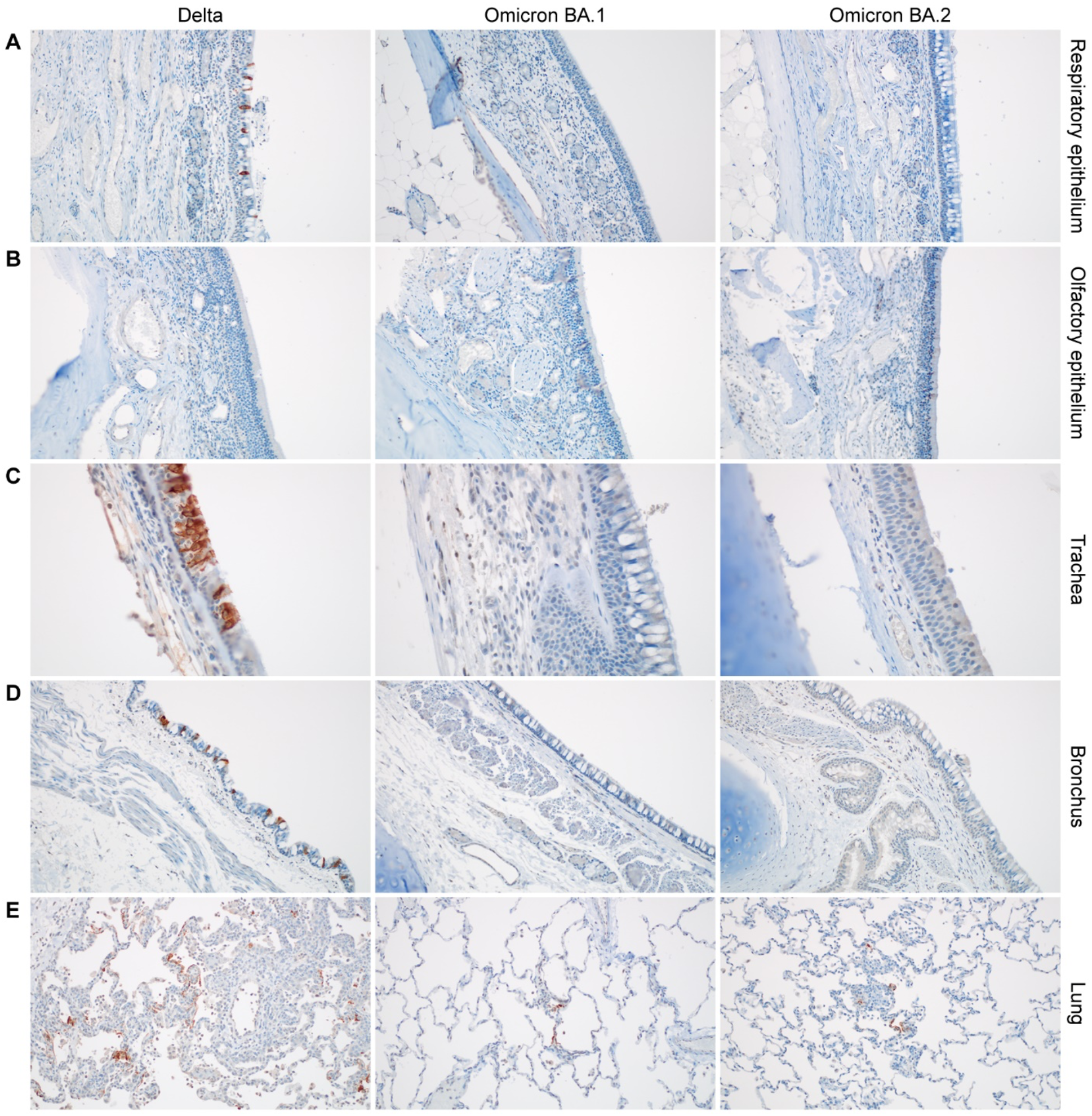
SARS-CoV-2 antigen staining in the respiratory tract of SARS-CoV-2 challenged rhesus macaques at 6 dpi. Serial sections of the samples described in Fig. 4. (A) Delta: Respiratory epithelial cell SARS-CoV-2 antigen staining (brown). Omicron BA.1: No SARS- CoV-2 antigen staining. Omicron BA.2: No SARS-CoV-2 antigen staining. (B) All: No SARS-CoV-2 antigen staining in olfactory epithelium (C) Delta: Tracheal mucosal SARS-CoV-2 antigen staining. Omicron BA.1: No SARS-CoV-2 antigen staining. Omicron BA.2: No SARS-CoV-2 antigen staining. (D) Delta: Bronchial mucosal SARS-CoV-2 antigen staining. Omicron BA.1: No SARS-CoV-2 antigen staining. Omicron BA.2: No SARS-CoV-2 antigen staining. (E) Delta: Multifocal and frequent SARS-CoV-2 antigen staining lining alveoli and intracellularly throughout the lung. Omicron BA.1: Example of extremely rare foci of immunoreactivity. Omicron BA.2: Example of extremely rare foci of immunoreactivity. Magnification A, B, D, E 200x; C 400x.

**Figure 6.**
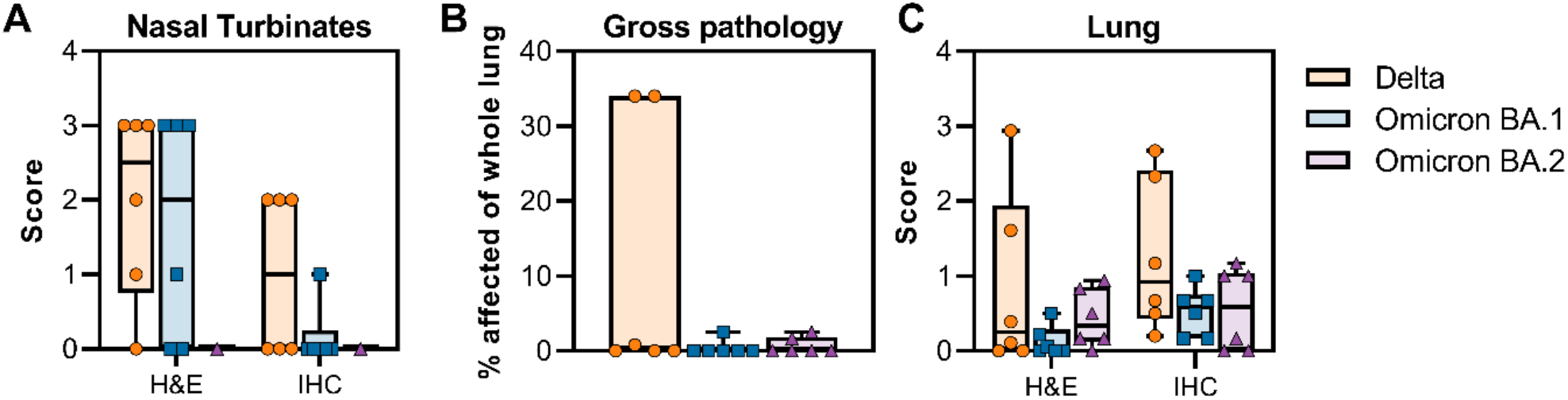
Scoring of pathology and SARS-CoV-2 antigen staining in nasal turbinate and lung tissue, and gross pathology in lung tissue. (A) Scoring between 0 (no pathology or staining) and 5 (severe pathology or diffuse staining) of nasal turbinates was done by a board- certified veterinary pathologist who was blinded to the study groups. (B) Gross pathology was scored per lung lobe (6 total), dorsal and ventral side. Percentage of the whole lung affected was then calculated. (C) Scoring between 0 (no pathology or staining) and 5 (severe pathology or diffuse staining) of lung tissue was done by a board-certified veterinary pathologist who was blinded to the study groups.

The trachea showed a milder inflammation than the nasal turbinates. Three out of six animals in the Delta group, all six Omicron BA.1 animals, and no Omicron BA.2 animals were found to have inflammation in the trachea. Surprisingly, only two animals had SARS-CoV-2 antigen, and both were challenged with the Delta variant. This may suggest that the virus was cleared from these antigen-negative tissues, or that inflammation was caused by repeated intubation of the animals (**Figure 4C, Figure 5C**).

Less inflammation was noted in the bronchi when compared to the trachea, two out of six Delta animals, three out of six Omicron BA.1 animals, and none of the Omicron BA.2 animals exhibited inflammation. Like the trachea, only two Delta challenged macaques had bronchial mucosal immunoreactivity to SARS-CoV-2 (**Figure 4D, Figure 5D**).

Gross lung lesions associated with SARS-CoV-2 pneumonia were identified as foci of consolidation and were noted in three animals in the Delta group, one animal in the Omicron BA.1 group, and two animals in the Omicron BA.2 group. The lung lesions in two animals in the Delta group affected a larger percentage of the total lung tissue (34%) than in the Omicron BA.1 and BA.2 groups (less than 3%) (**Figure S6B**). The observed features of SARS-CoV-2 pneumonia in this study included thickening of the alveolar septa with fibrin, edema and inflammatory cells, intra-alveolar inflammation, type II pneumocyte hyperplasia, reactive endothelial cells in blood vessels and perivascular inflammation. The inflammatory cells present included neutrophils, macrophages, and lymphocytes. When present, lesion severity ranged from minimal-to-mild in the Omicron BA.1 and BA.2 groups, and minimal-to-moderate in the Delta groups. Interestingly, two out of six animals in the Delta group developed lesions in much higher frequency and severity than the other four animals. SARS-CoV-2 antigen was rarely detected but present in type I pneumocytes and mononuclear cells in foci with and without features of pneumonia in all six Omicron BA.1 inoculated macaques and three out of six Omicron BA.2 challenged macaques. Comparatively, more SARS-CoV-2 antigen could be detected in all six of the Delta challenged group which ranged from rare to multifocal in severity (**Figure 4E, Figure 5E, Figure 6C**).

Overall, infection with all three VOCs resulted in lesions typical of SARS-CoV-2 pneumonia in macaques. Omicron BA.1 and BA.2 VOCs resulted in a lower number of lesions with lesser severity than observed in animals infected with the Delta VOC.

### Cytokines and chemokines are upregulated in animals inoculated with Delta

The presence of nine different cytokines was analyzed in nasosorption, serum, and BAL samples. Compared to baseline, nasosorption samples obtained from animals challenged with Delta showed an elevated immune response on all days. In particular, IL-1 receptor antagonist (IL1- RA), interleukin-6 (IL-6), IL-15, and tumor necrosis factor-α (TNF-α) were increased. In comparison, IL-6, IL-15 and TNF-α in nasosorption samples from animals inoculated with Omicron BA.1 and BA.2 were only moderately elevated or decreased compared to baseline samples. IL-1RA was elevated on 0-dpi in animals that received Omicron BA.1, and did not significantly increase over time (**Figure 7A, Figure S6**). Very few changes in cytokine and chemokine levels were observed in BAL samples: IL-1RA was upregulated in animals that were inoculated with Omicron BA.1 on 2-dpi (**Figure 7B, Figure S6**). In serum samples, similar responses were seen in all groups. IL-1RA was upregulated in all three groups on 2-dpi. TNF-α was slightly downregulated on 4- and 6-dpi, although the absolute values showed only a minor drop. On 6-dpi, the groups diverged slightly: IL-1RA, IL-6, and IL-8 were upregulated in the Delta group, whereas IL-6 and MCP-1 were downregulated in the Omicron BA.1 and BA.2 groups (**Figure 7C, Figure S6)**.

**Figure 7.**
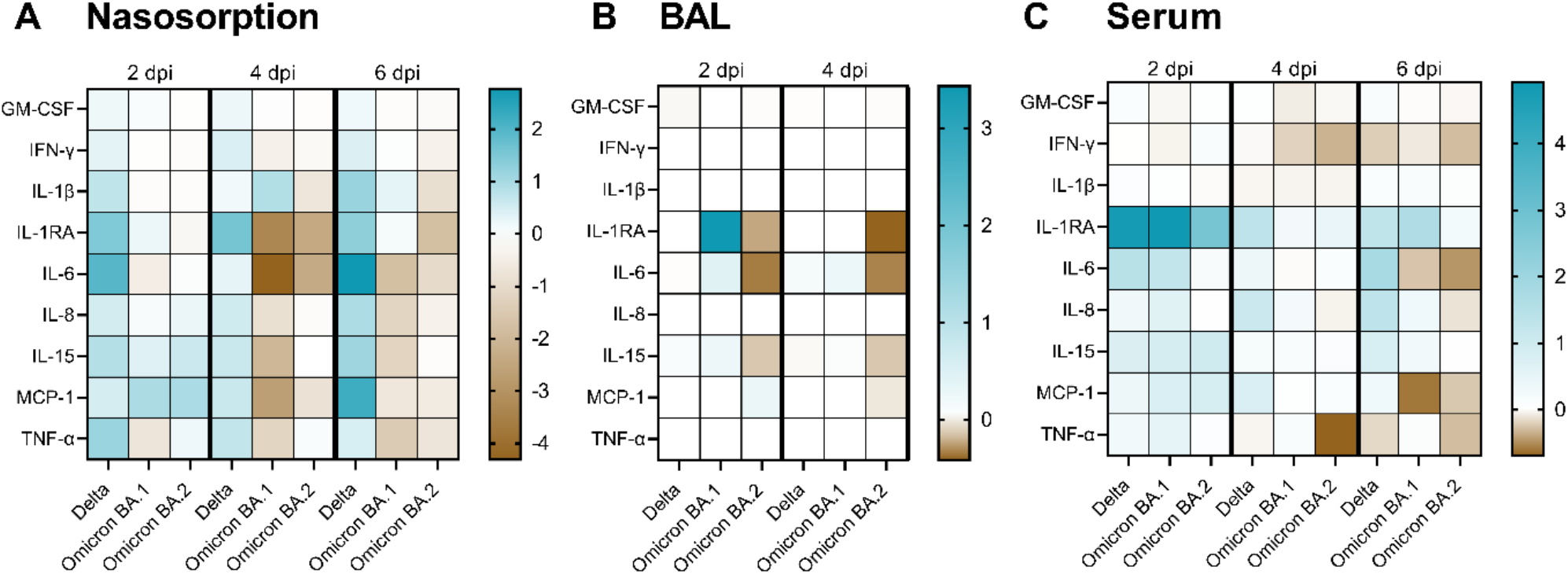
Cytokine and chemokines in nasosorption, BAL, and serum samples were downregulated in animals inoculated with Omicron VOCs compared to Delta VOC. Cytokine and chemokine levels were determined in nasosorption (A), BAL (B) and serum (C) samples obtained at pre-challenge, 2, 4, and 6 (nasosorption and serum only) days post-challenge. Fold-changes were calculated over baseline (pre-challenge values) and median log2 values are displayed.

## Discussion

Severity of disease is an important variable when considering a public health response, more so when the infectious agent causing disease has become as wide-spread as SARS-CoV-2. Omicron is the first VOC which has been reported to cause less severe disease in the human population than the preceding VOC wave (8,9). Disease severity is likely to be reduced by the presence of SARS-CoV-2 specific immunity in the population, either through vaccination or previous infections (10). In South Africa, where Omicron rapidly displaced the Delta VOC, a lower proportion of reported infections ended in hospitalizations and deaths during the Omicron wave as compared to previous waves with the ancestral, Beta, and Delta variants (11). However, the seroprevalence of SARS-CoV-2 IgG was determined to be 68.4% before the Omicron wave, compared to 19.1% after the Beta wave. The increased rates of immunity generated either by vaccine or infection likely plays a crucial role in the reduction of disease severity (12). Whether disease severity would have been reduced in the absence of pre-existing immunity is currently not known.

Studies utilizing both hamsters and mice have shown that infection with Omicron BA.1 resulted in a lack of weight loss and lower viral burdens in the upper and lower respiratory tract compared to other SARS-CoV-2 VOCs (13). Furthermore, viral loads in nasal swabs obtained from NHPs inoculated with Omicron BA.1 appear low compared to viral loads in nasal swabs obtained from NHPs inoculated with a Lineage A isolate, whereas viral load in BAL appears similar (14,15).

Here, we show that rhesus macaques infected with Omicron BA.1 or BA.2 behave very similarly. Animals inoculated with Omicron BA.1 or BA.2 shed less virus and have a lower virus load in the lower respiratory tract than rhesus macaques infected with the Delta variant. This is accompanied by a reduction in observed clinical signs of disease, inflammatory lesions in the respiratory tract, and a decrease in the innate immune response in Omicron-inoculated animals compared to Delta-inoculated animals. Whereas the detection of viral RNA was mostly limited to the respiratory tract in Omicron-inoculated animals, viral RNA in Delta-inoculated animals was found in extra-respiratory tissues. Overall, these results support the notion that Omicron infection results in less severe disease, even in the absence of pre-existing immunity. It is possible that this difference is driven by the S protein of Omicron. In our entry studies, we show a reduced entry of Omicron compared to Delta for both human and rhesus macaque ACE2 in a BHK cell line. A similar difference in entry has been observed in Calu-3 and A549 cell lines, but not HEK cell lines, which may be driven by the TMPRSS2-independent, cathepsin-dependent endosomal entry pathway that Omicron favors compared to Delta (16).

The reduction in shedding of viral RNA we observed in animals inoculated with Omicron compared to Delta aligns with some of the shedding data published on vaccinated and unvaccinated individuals (10), although not all (17). Puhach *et al*. determined viral load and infectious virus titers in nasopharyngeal samples of 384 symptomatic individuals and did not find a difference in viral load or infectious virus (17). Chaguza *et al*. analyzed 37,877 nasal swab samples and showed consistently lower viral loads for samples obtained from participants infected with Omicron compared to Delta, independent of vaccination status (10). Since the difference in cycle threshold (Ct) values are subtle in this study (less than 1 cycle), it is possible that in humans, the sample numbers must be high to show significant differences in virus loads. We compared the amount of virus shed and detected in respiratory tract tissue between all six VOCs. Previously we showed there was no difference in the amount of viral RNA shed in nose swabs, BAL, or BCB samples for D614G, Alpha, and Beta VOCs. Interestingly, in nasal swabs, Omicron BA.1 and BA.2 were most like D614G, Alpha, and Beta, whereas the viral load detected in nasal swabs from animals inoculated with the Delta VOC was higher. In BCB samples, viral load was again highest for the Delta VOC, whereas no differences between the VOCs were observed in BAL samples. In lung tissue, the amount of viral RNA was significantly higher for animals inoculated with the Delta VOC when compared to all other VOCs. Thus, Delta is the VOC most efficient at replication in the naïve rhesus macaque model.

Nonetheless, the Omicron VOC has replaced the Delta VOC in the human population, and our study was not designed to address this question. Omicron is antigenically the most distant VOC (18) and recent studies suggest that Omicron variants can readily overcome immunity acquired from previous infection with earlier variants and vaccination (19–23). The rise in Omicron cases could be a combination of immune evasion, waning immunity, relaxation of COVID-19 restrictions, and other factors that may affect transmission, such as reduced symptoms caused by Omicron resulting in prolonged contact with other humans. None of these features were investigated in our study, and their influence can thus not be assessed.

We assessed the cytokine and chemokine response in three different samples: nasosorption samples represent the upper respiratory tract, BAL samples represent the lower respiratory tract, and serum samples represent the systemic response. In the upper respiratory tract, cytokines and chemokines were upregulated to higher levels in Delta-inoculated animals than in Omicron BA.1 or BA.2-inoculated animals, whereas the systemic response was comparable. This is likely directly correlated to the amount of antigen: we consistently found higher viral loads in nasal swabs, BCBs and lung tissues of animals inoculated with Delta compared to Omicron. This highlights the need for obtaining samples from the site of virus replication to obtain a full understanding of the innate immune response, both in animal studies and in patients.

Here we show that in naïve rhesus macaques the Delta VOC replicated to higher viral loads than the D614G, Alpha, Beta, and Omicron BA.1 and BA.2 variants, resulting in more virus shed and increased replication in lung tissue. Although similar results were found in small animal models of SARS-CoV-2 infection, this study was the first to directly compare Delta, Omicron BA.1, and Omicron BA.2 in a species that share the same ACE2 receptor sequences to humans. The reduction in viral load, disease and pathology detected following Omicron BA.1 and BA.2 infection is reflective of what is seen in the human population. Finally, this study further validates the rhesus macaque model for continued evaluation and comparison of the phenotype and pathogenicity of novel emerging variants.

## Materials and Methods

### Study Design

Three groups of six rhesus macaques were inoculated with either SARS-CoV-2 VOC Delta AY.106 (hCoV-19/USA/MD-HP05647/2021, EPI_ISL_2331496), SARS-CoV-2 VOC Omicron BA.1 (hCoV-19/USA/GA-EHC-2811C/2021, EPI_ISL_7171744), or SARS-CoV-2 VOC Omicron BA.2 (hCoV-19/Japan/UT-NCD1288-2N/2022, EPI_ISL_9595604). Eighteen rhesus macaques between the ages of 2 and 22 were randomly divided into groups of six animals consisting of three females and three males. The age range of each group were as follows; Delta was 3-22 years, BA.1 was 6-19 years and BA.2 was 2-4years. Each group of animals was housed in a separate room. The animals were inoculated as previously described (1). Briefly, NHPs were inoculated intranasally (0.5 mL) and intratracheally (4 mL) with a total dose of 2 × 10^6^ TCID_50_ virus dilution in sterile Dulbecco’s modified Eagle’s medium (DMEM). The inoculum dose was confirmed by titration on Vero E6 cells. The same person, blinded to the study groups, assessed the animals throughout the study using a standardized scoring sheet (24) and based on the evaluation of the following criteria: general appearance and activity, appearance of skin and coat, discharge, respiration, feces and urine output, and appetite. Area under the curve analysis was performed using Graphpad Prism 9.3.1 to obtain a single clinical score value per animal. Clinical exams were performed on 0-, 2-, 4-, and 6-dpi. Swabs (nose, throat, and rectal), nasosorption samples, bronchial cytology brush samples, and blood were collected at all exam dates.

Nasosorption samples were collected as previously described (25). On -10-, 2- and 4-dpi, animals were intubated and BALs were performed using 10 mL of sterile saline. Ventrodorsal and right/left lateral thoracic radiographs were taken before any other procedures. Two board-certified clinical veterinarians blinded to study groups scored the radiographs for the presence of pulmonary infiltrates according to a standard scoring system as previously described (1). Scores may range from 0 to 18 for each animal on each exam day. On 6-dpi, all animals were euthanized; after euthanasia, necropsies were performed and 27 tissue samples were collected.

### Ethics and Biosafety

The Institutional Animal Care and Use Committee (IACUC) of Rocky Mountain Laboratories, National Institutes of Health (NIH) approved all animal experiments. Experiments are carried out in an Association for Assessment and Accreditation of Laboratory Animal Care (AAALAC) International–accredited facility, according to the institution’s guidelines for animal use, following the guidelines and basic principles in the NIH *Guide for the Care and Use of Laboratory Animals*, the Animal Welfare Act, U.S. Department of Agriculture, and the U.S. Public Health Service Policy on Humane Care and Use of Laboratory Animals. Rhesus macaques were single-housed in adjacent primate cages, which allow social interactions. The animal room was climate-controlled with a fixed light-dark cycle (12-hour light/12-hour dark). Commercial monkey chow was provided twice daily. Water was available ad libitum. The diet was supplemented with treats, vegetables, or fruit at least once a day. Environmental enrichment consisted of a variety of human interaction, manipulanda, commercial toys, videos, and music. Animals were monitored at least twice daily throughout the experiment. The Institutional Biosafety Committee (IBC) approved work with SARS-CoV-2 under Biosafety Level 3 conditions as well as subsequent sample inactivation for removal of specimens from high containment (26).

### Virus and cells

In this study, three SARS-CoV-2 strains were utilized: Delta AY.106 VOC (hCoV-19/USA/MD- HP05647/2021, EPI_ISL_2331496) was obtained from Andrew Pekosz, Johns Hopkins Bloomberg School of Public Health; the Omicron BA.1 VOC (hCoV-19/USA/GA-EHC- 2811C/2021, EPI_ISL_7171744) was obtained from Mehul Suthar, Emory University School of Medicine, and the Omicron BA.2 VOC (hCoV-19/Japan/UT-NCD1288-2N/2022, EPI_ISL_9595604) was obtained from Peter Halfmann, University of Wisconsin. VeroE6 cells (provided by Professor Ralph Baric, University of North Carolina at Chapel Hill) were maintained in DMEM supplemented with 10% fetal bovine serum, 1 mM L-glutamine, penicillin (50 U/mL), and streptomycin (50 μg/mL; DMEM10). Mycoplasma testing was performed monthly, with no mycoplasma detected in cells or stocks used in this study.

All virus propagation was performed in VeroE6 cells in DMEM2 (DMEM supplemented with 2% fetal bovine serum, 1 mM L-glutamine, penicillin (50 U/mL), and streptomycin (50 μg/mL)). Sequencing confirmed there were no mutations in the consensus of the Delta and Omicron BA.1 strains. The Omicron BA.2 strain had an A116V substitution in NSP16 in 69% of the reads.

### SARS-CoV-2 entry in BHK cells using human and rhesus macaque ACE2 receptors

BHK cells were seeded in black 96-well plates at 6.0 × 10^5^ cells/mL one day prior to transfection (n = 8 wells/variant, experiment repeated twice). The next day, cells were transfected with 100 ng of human or rhesus ACE2 receptor plasmid DNA using polyethylenimine (Polysciences). After 24 h, cells were inoculated with 100 μL of pseudotype stocks at a 1:10 dilution. Plates were then centrifuged at 1200 × g at 4 °C for 1 h and incubated overnight at 37 °C. Approximately 16–20 h post-infection, Bright-Glo luciferase reagent (Promega) was added to each well, at a 1:1 dilution, and luciferase was measured. Relative entry was calculated by normalizing the relative light unit for variant S pseudotypes to the plate relative light unit average for the lineage A spike pseudotype.

### Plasmids

Plasmids of the human and rhesus macaque ACE2 receptors and S coding sequences for SARS-CoV-2 lineage A, Delta, Omicron BA.1, and Omicron BA.2 were developed. All plasmids used the pcDNA3.1^+^ vector (GenScript) and were verified by Sanger sequencing (ACGT). Because coronavirus S proteins with a 19 aa deletion at the C-terminus have previously been found to have an increase in incorporation for virions of VSV (27), all S sequences in the plasmids included the 19 aa truncation. Additionally, the S sequences were codon-optimized for human cells as well as appended with a 5′ Kozak expression sequence (GCCACC) and 3′ tetra-glycine linker followed by nucleotides encoding a FLAG-tag sequence (DYKDDDDK).

### Pseudotype production

Pseudotype production followed a previously established protocol (28). Briefly, plates pre-coated with poly-L-lysine (Sigma–Aldrich) were seeded with 293T cells and transfected the following day with 1,200 ng of empty plasmid and 400 ng of plasmid encoding coronavirus S or no-S plasmid control (green fluorescent protein (GFP)). After 24 h, transfected cells were infected with VSVΔG seed particles pseudotyped with VSV-G. After an hour of incubating with intermittent shaking at 37 °C, cells were washed four times and incubated in 2 mL DMEM2 for 48 h. Supernatants were collected, centrifuged at 500xg for 5 min, aliquoted, and stored at −80 °C.

### Virus RNA extraction and quantitative polymerase chain reaction

RNA was extracted from liquid samples using a QiaAmp Viral RNA kit (Qiagen) according to the manufacturer’s instructions, whereas tissue was homogenized and extracted using the RNeasy kit (Qiagen) according to the manufacturer’s instructions. Viral gRNA (29) and sgRNA (30) were detected using specific assays: RNA (5 μl) was tested with the QuantStudio (Thermo Fisher Scientific) according to instructions of the manufacturer. SARS-CoV-2 standards with known genome copies were run in parallel to allow for quantification.

### Histopathology

Tissues were fixed for a minimum of 7 days in 10% neutral-buffered formalin and embedded in paraffin, followed by staining with hematoxylin and eosin, or using a custom-made rabbit antiserum against SARS-CoV-2 N at a 1:1000 dilution. Stained slides were analyzed by a board-certified veterinary pathologist who was blinded to the study groups. Histologic lesion severity was scored per lung lobe according to a standardized scoring system evaluating the presence of interstitial pneumonia, type II pneumocyte hyperplasia, edema and fibrin, and perivascular lymphoid cuffing as follows: 0, no lesions; 1, minimal (1 to 10% of lobe affected); 2, mild (11 to 25%); 3, moderate (26 to 50%); 4, marked (51 to 75%); and 5, severe (76 to 100%). Presence of viral antigen was scored per lung lobe according to a standardized scoring system: 0, none; 1, rare/few; 2, scattered; 3, moderate; 4, numerous; and 5, diffuse.

### Cytokine and chemokine analysis

The U-PLEX Biomarker Group 1 (NHP) Assay kit (MSD, K15068L-2) from MSD was used to test the presence of nine cytokines (GM-CSF, IFN-γ, IL-1β, IL-1RA, IL-6, IL-8, IL-15, MCP-1, and TNF-α) in nasosorption, serum, and BAL NHP samples. The plates were immediately read using the Meso Quickplex instrument (MSD, K15203D). The data was extracted from the plates using the MSD Workbench 4.0 software. The fold-change compared to pre-challenge samples and Log_2_ values for the BAL, serum and nasosorption samples were calculated using Microsoft Excel, and graphed using GraphPad Prism 9.1.1 (225) software.

### Statistical analysis

Statistical analyses were performed using GraphPad Prism software version 8.2.1. For all analyses, a P value of 0.05 was used as cutoff for statistical significance.

## Acknowledgements

We thank Tina Thomas, Rebecca Rosenke, and Dan Long for assistance with histology. Brandon Bailes, Richard Cole, Lydia Crawford, Lisa Heaney, Corey Henderson, Taylor Lippincott, Travis Spencer, Kathy Cordova and Marissa Woods of the RMVB animal care staff. Meaghan Flagg, Myndi Holbrook, Greg Saturday, Craig Martens, Patrick Hanley, Kent Barbian, Stacey Ricklefs, Sarah Anzick, and Dana Scott for technical assistance; Anita Mora for assistance with the figures; and Andrew Pekosz, Mehul Suthar, Peter Halfmann for assistance in acquiring the VOC isolates.

## Author contributions

Conceptualization: N.v.D., H.F., V.J.M., E.d.W., and K.R.; investigation: N.v.D., M.S., T.A.S., C.K.Y., L.P.P, F.B., Z.A.W., M.L., J.E.S., B.N.W., K.M.W, S.G., F.F., J.L., C.S., and K.R.; writing (original draft): N.v.D, M.S., and K.R.; writing (review and editing): all authors. Funding: V.J.M. and H.F.

## Data and materials availability

All data included in this manuscript have been deposited in Figshare at XXX.

---

**Table S1.**
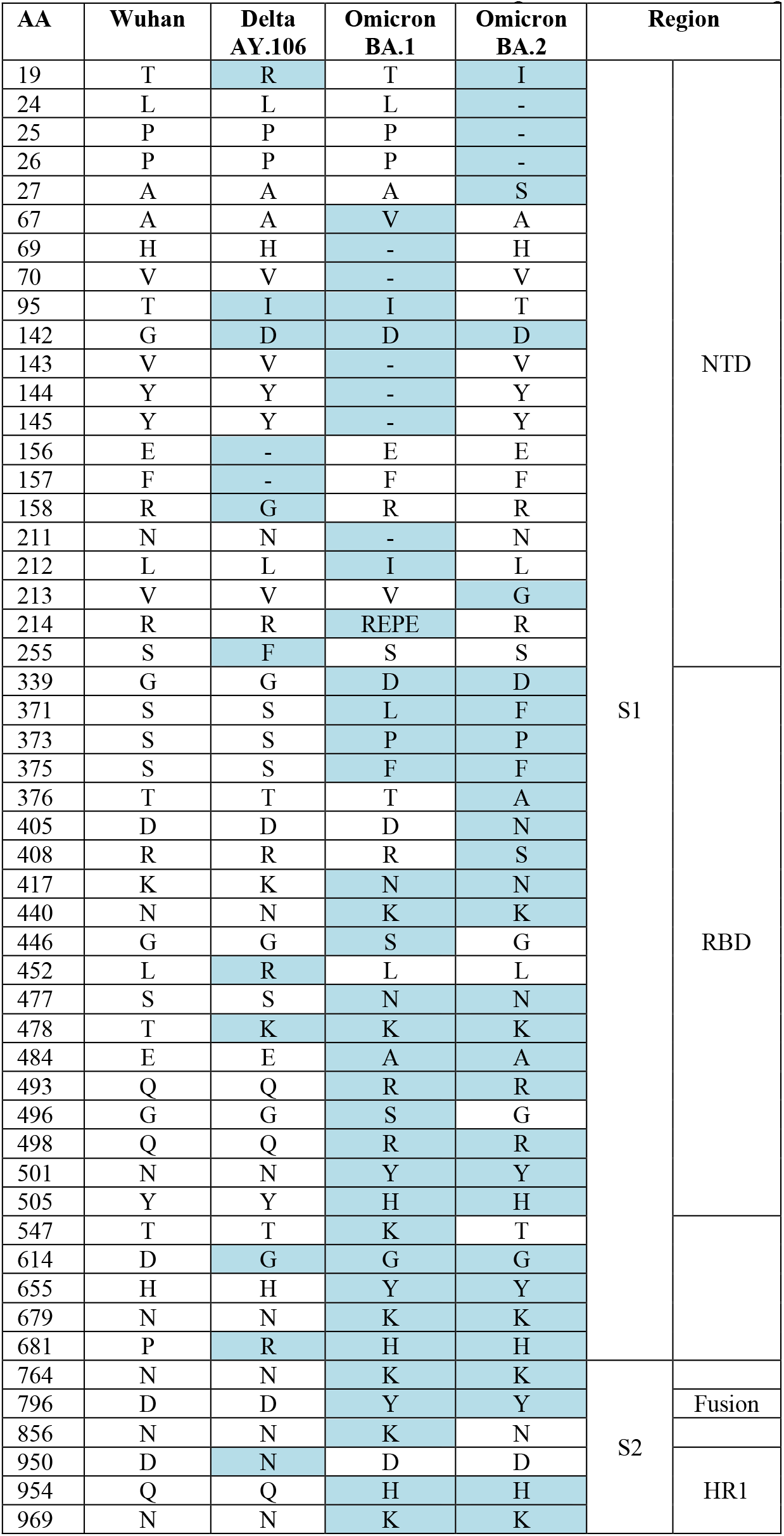
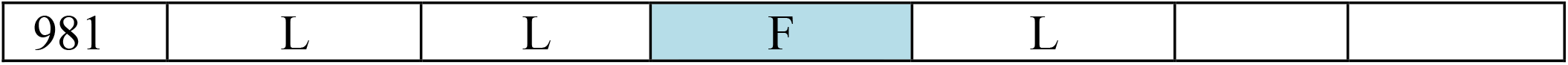
NHPs were inoculated with SARS-CoV-2 VOCs Delta AY.106 (hCoV-19/USA/MD- HP05647/2021, EPI_ISL_2331496), Omicron BA.1 (hCoV-19/USA/GA-EHC-2811C/2021, EPI_ISL_7171744), or Omicron BA.2 (hCoV-19/Japan/UT-NCD1288-2N/2022, EPI_ISL_9595604). No substitutions in the S protein compared to published sequence were found. All amino acid substitutions compared to ancestral S protein Wuhan are detailed below.

**Figure S1.**
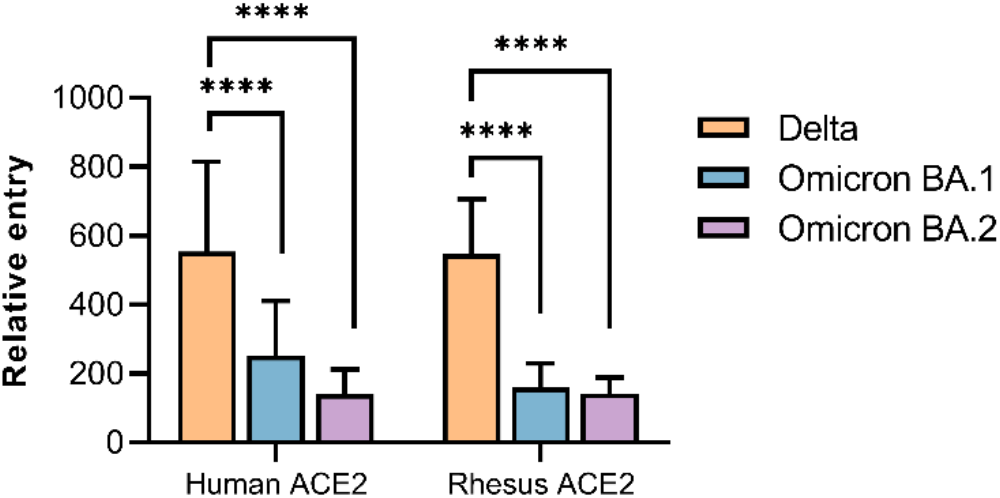
Comparison of entry of SARS-CoV-2 S proteins to human and rhesus macaque ACE2. BHK cells were transfected with either human ACE2 or rhesus ACE2 and subsequently infected with pseudotyped VSV reporter particles with the S proteins of Delta, Omicron BA.1, or Omicron BA.2. Luciferase expression was measured, and relative entry of the VOCs was calculated over no spike pseudotype. N=16, combined from two separate experiments. Statistical analysis was performed using a one-way ANOVA with Tukey’s multiple comparisons test.

**Figure S2.**
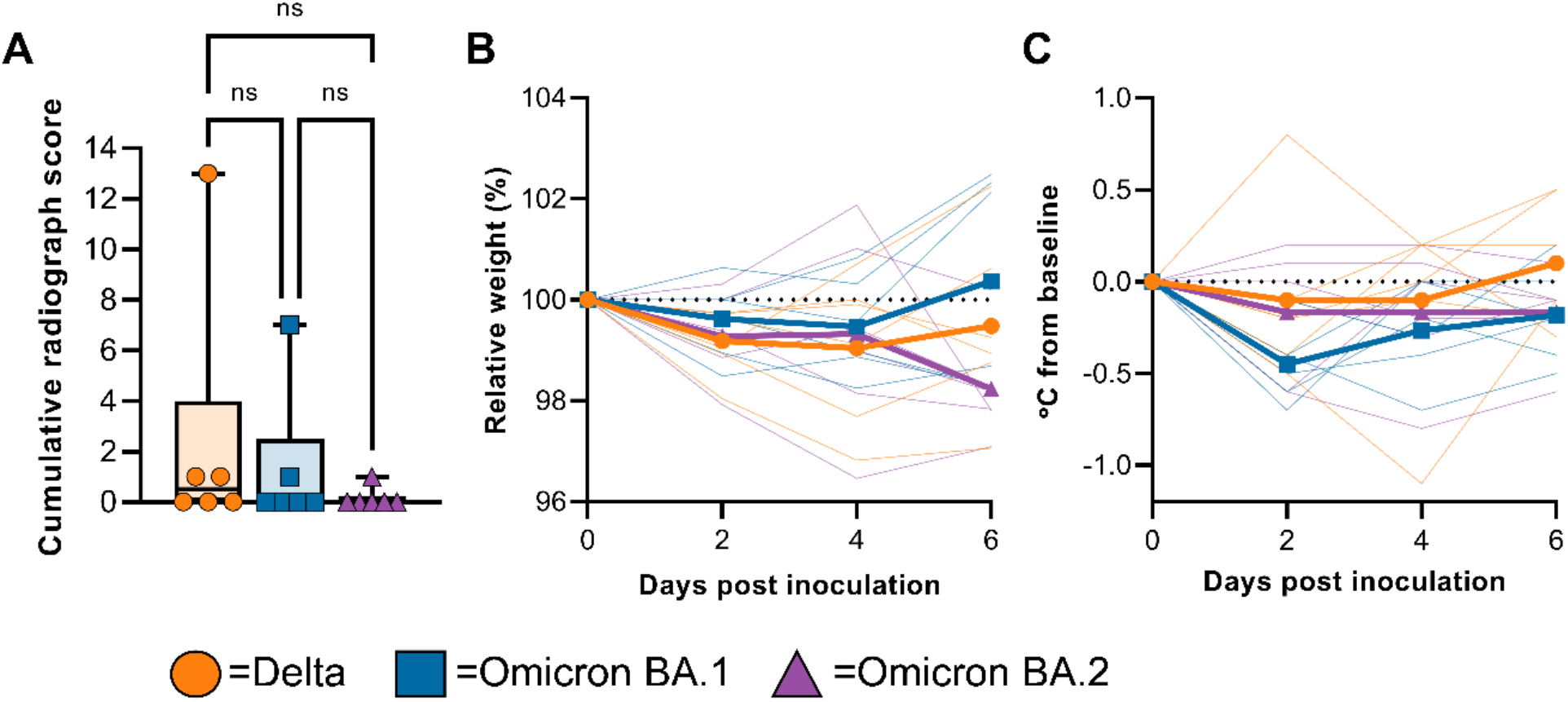
Limited signs of disease on radiographs, weight and temperature. (A) Minimum- to-maximum boxplot of ventrodorsal radiographs taken on exam days. Individual lobes were scored by a clinical veterinarian according to a standardized scoring system and totaled. Statistical analysis was performed using a Kruskal-Wallis test with Dunn’s multiple comparisons. (B) The relative weight compared to the day of challenge (dotted line) is shown per group (median, thick line) as well as per individual (thin lines). (C) Body temperature is indicated as deviation from baseline at the day of challenge (dotted line) and shown per group (median, thick line) as well as per individual (thin lines).

**Figure S3.**
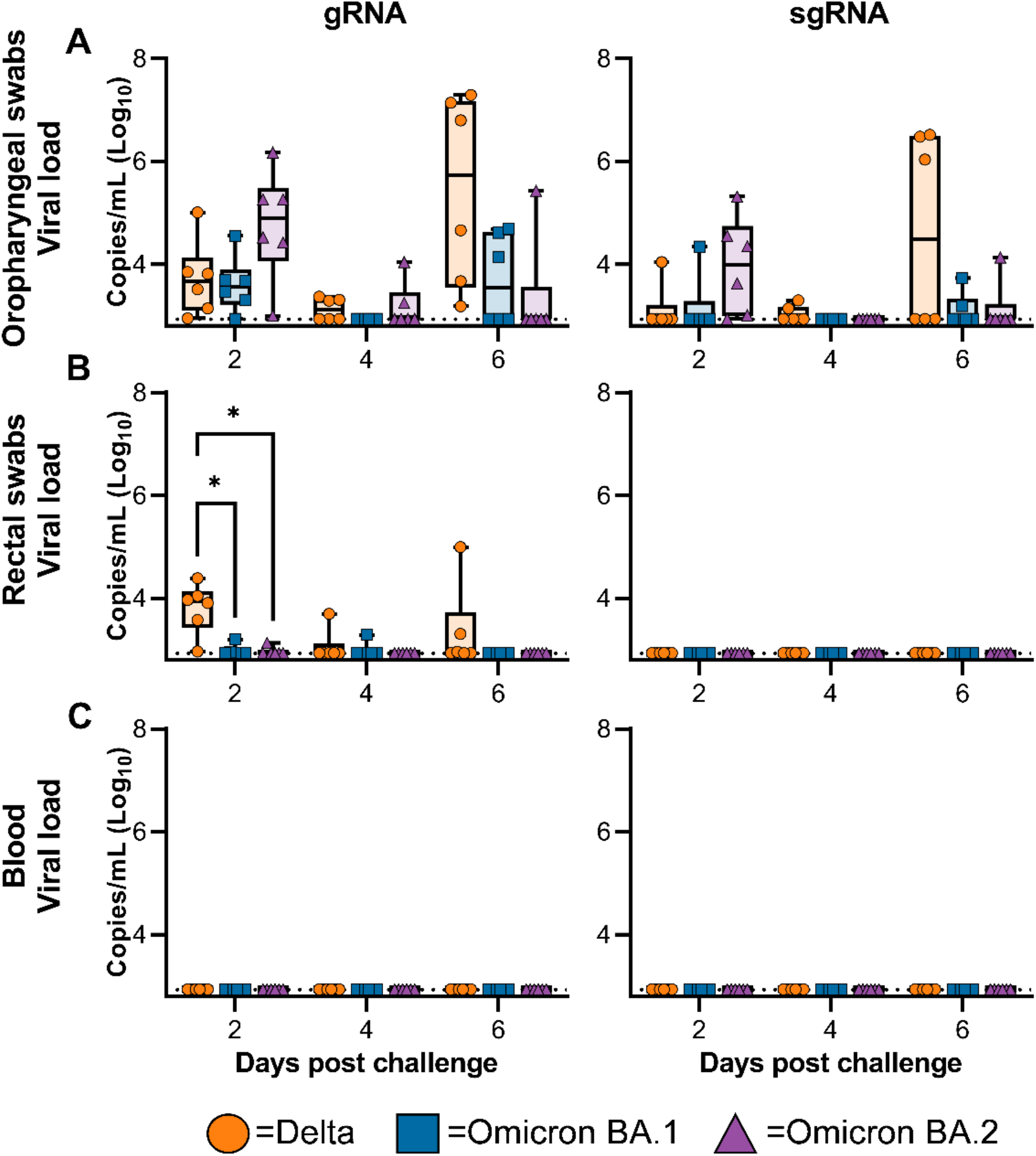
Limited differences in shedding of viral RNA were detected in throat swabs, rectal swabs, and blood. Boxplot (minimum-to-maximum) of viral gRNA (left panel) and sgRNA (right panel) in throat swabs (A), rectal swabs (B), and blood (C) taken on 2-, 4-, and 6- dpi. Statistical significance was determined via a two-way ANOVA with the Geisser-Greenhouse correction followed by the Tukey test for multiple comparisons.

**Figure S4.**
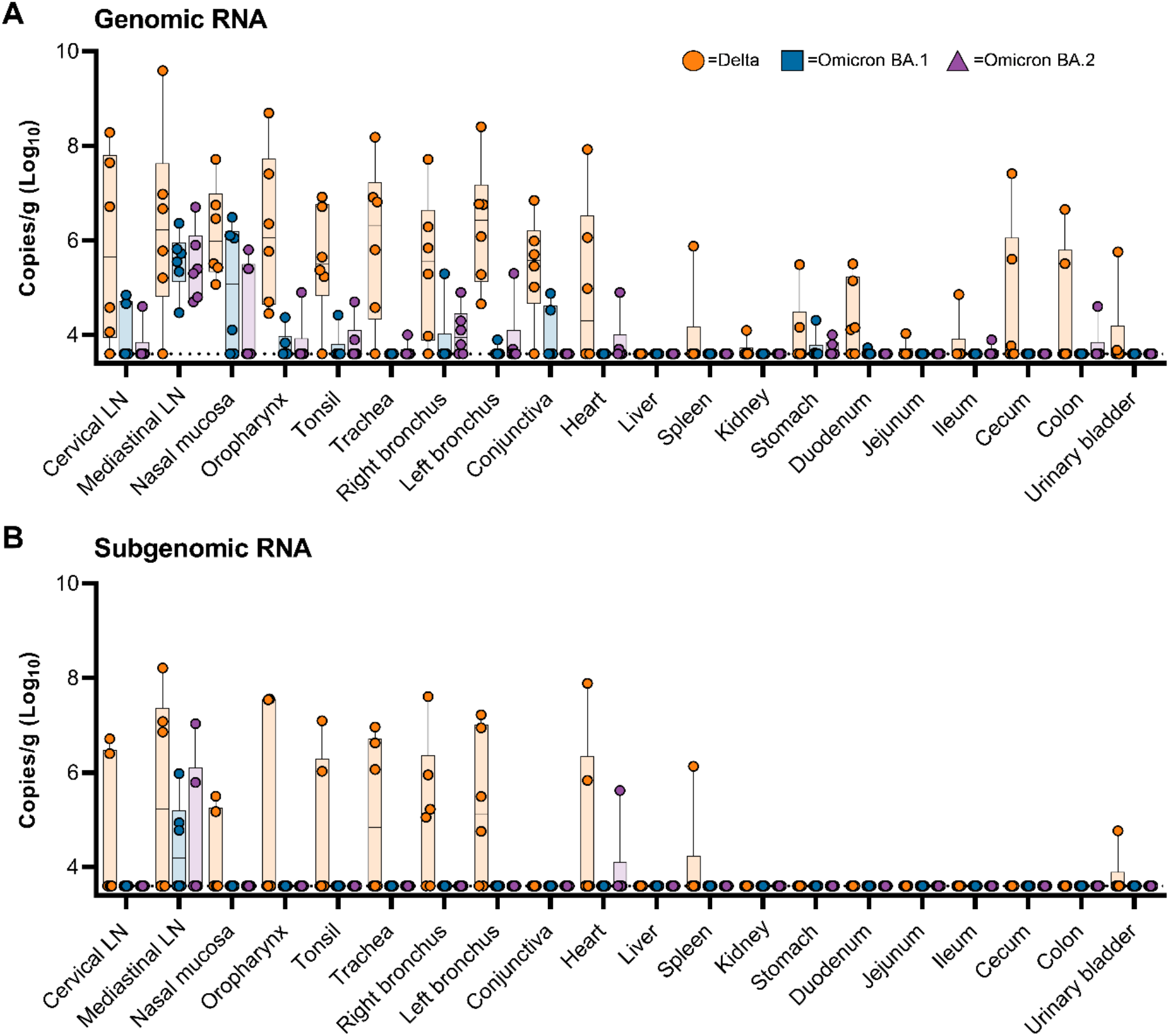
Viral loads in non-respiratory tissues is limited for Omicron BA.1 and BA.2 inoculated animals on 6-dpi. (A) Boxplot (minimum-to-maximum) of gRNA detected in tissues. (B) Boxplot (minimum-to-maximum) of sgRNA detected in tissues.

**Figure S5.**
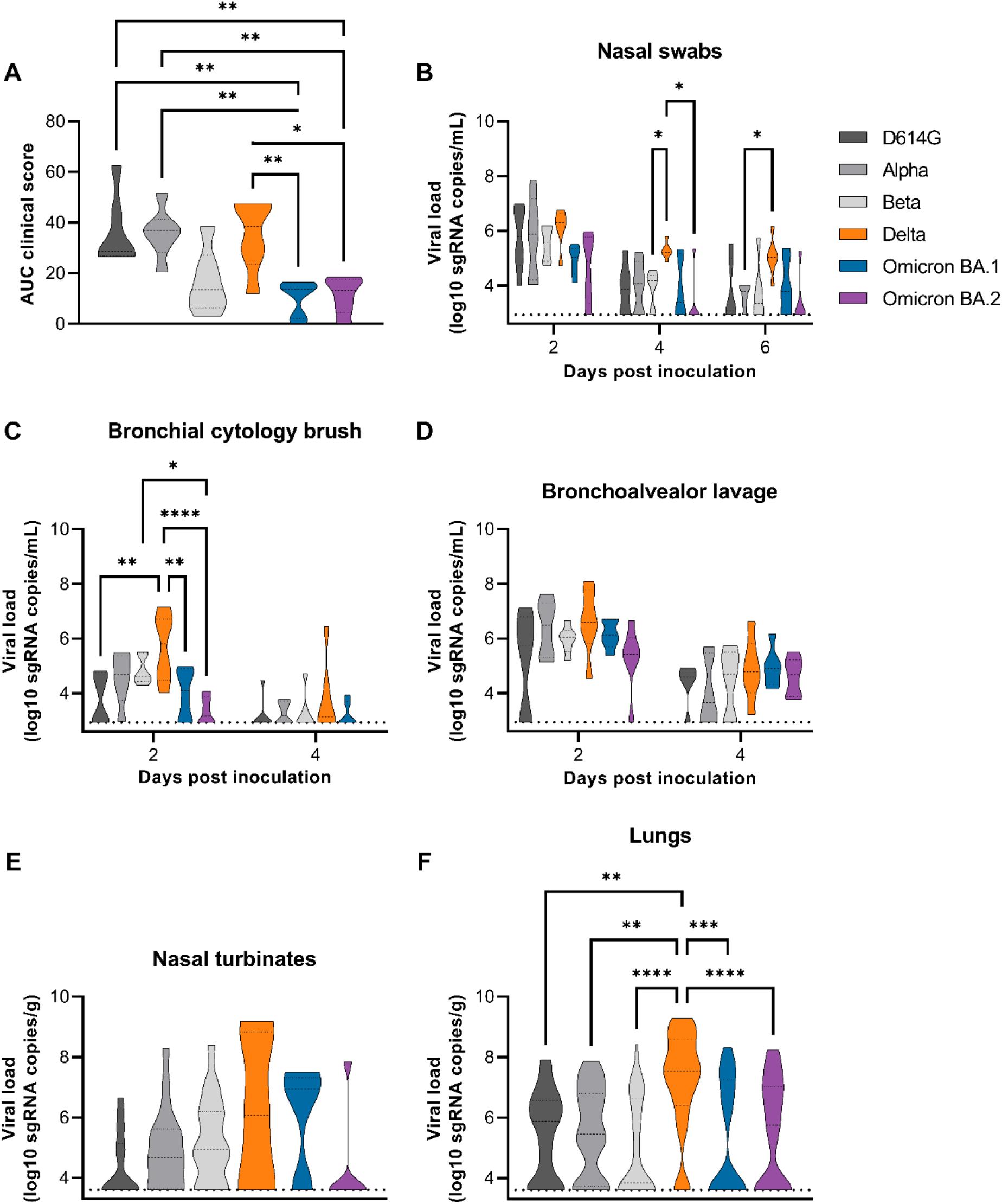
Clinical score and viral load comparison in respiratory tract samples between D614G, Alpha, Beta, Delta, Omicron BA.1, and Omicron BA.2 variants. Truncated violin plot of clinical score (A), viral load in nasal swabs (B), BCBs (C), BAL fluid (D), nasal turbinate tissue (E), and lung tissue (F). Samples in grey are from a previously published study (7). Dotted line = qualitative limit of detection (10 copies per reaction). Statistical analyses done via ordinary one-way ANOVA followed by Holm- Šídák’s multiple comparisons test (A), two-way ANOVA followed by multiple comparison via Tukey (B, C, D), Kruskal-Wallis test followed by multiple comparison via Dunn’s (D, E).

**Figure S6.**
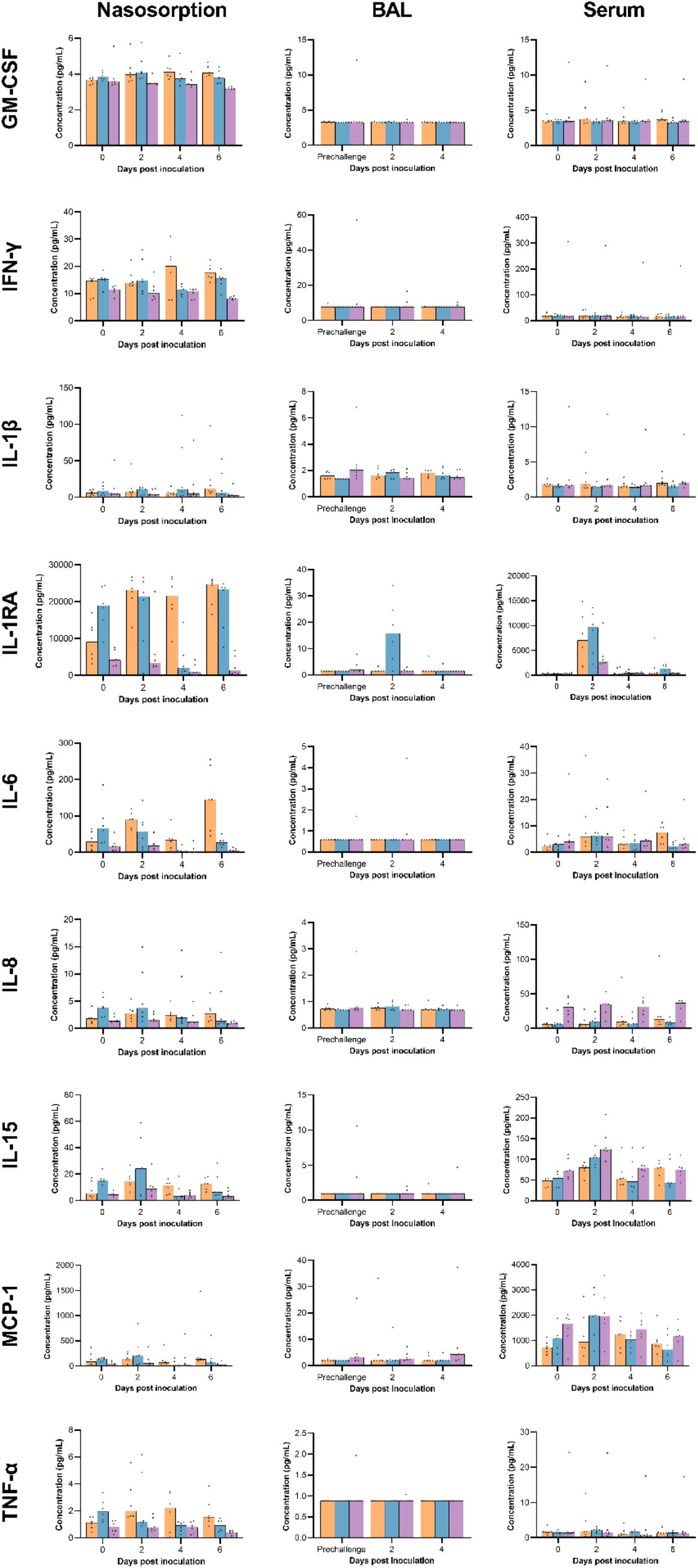
Absolute values of cytokines and chemokines measured in nasosorption, BAL, and serum samples. Bar graphs of median and individual values. Orange = Delta; Blue = Omicron BA.1; Purple = Omicron BA.2

## References

1. Konings F, Perkins MD, Kuhn JH, Pallen MJ, Alm EJ, Archer BN, et al. SARS-CoV-2 Variants of Interest and Concern naming scheme conducive for global discourse. Nat Microbiol. 2021 Jul;6(7):821–3.

2. WHO. COVID-19 Weekly Epidemiological Update, 25 February 2021 [Internet]. Available from: https://www.who.int/docs/default-source/coronaviruse/situation-reports/20210225_weekly_epi_update_voc-special-edition.pdf

3. Tao K, Tzou PL, Nouhin J, Gupta RK, de Oliveira T, Kosakovsky Pond SL, et al. The biological and clinical significance of emerging SARS-CoV-2 variants. Nat Rev Genet. 2021 Dec;22(12):757–73.

4. Riley S, Wang H, Eales O, Haw D, Walters CE, Ainslie KEC, et al. REACT-1 round 12 report: resurgence of SARS-CoV-2 infections in England associated with increased frequency of the Delta variant [Internet]. Infectious Diseases (except HIV/AIDS); 2021 Jun [cited 2022 Mar 23]. Available from: http://medrxiv.org/lookup/doi/10.1101/2021.06.17.21259103

5. Bolze A, Luo S, White S, Cirulli ET, Wyman D, Dei Rossi A, et al. SARS-CoV-2 variant Delta rapidly displaced variant Alpha in the United States and led to higher viral loads. Cell Reports Medicine. 2022 Mar;3(3):100564.

6. Nikolaidis M, Papakyriakou A, Chlichlia K, Markoulatos P, Oliver SG, Amoutzias GD. Comparative Analysis of SARS-CoV-2 Variants of Concern, Including Omicron, Highlights Their Common and Distinctive Amino Acid Substitution Patterns, Especially at the Spike ORF. Viruses. 2022 Mar 29;14(4):707.

7. Munster VJ, Flagg M, Singh M, Yinda CK, Williamson BN, Feldmann F, et al. Subtle differences in the pathogenicity of SARS-CoV-2 variants of concern B.1.1.7 and B.1.351 in rhesus macaques. Sci Adv. 2021 Oct 22;7(43):eabj3627.

8. Sigal A, Milo R, Jassat W. Estimating disease severity of Omicron and Delta SARS-CoV-2 infections. Nat Rev Immunol. 2022 May;22(5):267–9.

9. Wolter N, Jassat W, Walaza S, Welch R, Moultrie H, Groome M, et al. Early assessment of the clinical severity of the SARS-CoV-2 omicron variant in South Africa: a data linkage study. The Lancet. 2022 Jan;399(10323):437–46.

10. Chaguza C, Coppi A, Earnest R, Ferguson D, Kerantzas N, Warner F, et al. Rapid emergence of SARS-CoV-2 Omicron variant is associated with an infection advantage over Delta in vaccinated persons. Med. 2022 Apr;S2666634022001386.

11. Madhi SA, Kwatra G, Myers JE, Jassat W, Dhar N, Mukendi CK, et al. Population Immunity and Covid-19 Severity with Omicron Variant in South Africa. N Engl J Med. 2022 Apr 7;386(14):1314–26.

12. Mutevedzi PC, Kawonga M, Kwatra G, Moultrie A, Baillie V, Mabena N, et al. Estimated SARS-CoV-2 infection rate and fatality risk in Gauteng Province, South Africa: a population-based seroepidemiological survey. International Journal of Epidemiology. 2022 May 9;51(2):404–17.

13. Halfmann PJ, Iida S, Iwatsuki-Horimoto K, Maemura T, Kiso M, Scheaffer SM, et al. SARS-CoV-2 Omicron virus causes attenuated disease in mice and hamsters. Nature. 2022 Mar 24;603(7902):687–92.

14. Gagne M, Moliva JI, Foulds KE, Andrew SF, Flynn BJ, Werner AP, et al. mRNA-1273 or mRNA-Omicron boost in vaccinated macaques elicits comparable B cell expansion, neutralizing antibodies and protection against Omicron [Internet]. Immunology; 2022 Feb [cited 2022 Feb 8]. Available from: http://biorxiv.org/lookup/doi/10.1101/2022.02.03.479037

15. Corbett KS, Nason MC, Flach B, Gagne M, O’Connell S, Johnston TS, et al. Immune correlates of protection by mRNA-1273 vaccine against SARS-CoV-2 in nonhuman primates. Science. 2021 Sep 17;373(6561):eabj0299.

16. Willett BJ, Grove J, MacLean OA, Wilkie C, De Lorenzo G, Furnon W, et al. SARS-CoV-2 Omicron is an immune escape variant with an altered cell entry pathway. Nat Microbiol [Internet]. 2022 Jul 7 [cited 2022 Jul 11]; Available from: https://www.nature.com/articles/s41564-022-01143-7

17. Puhach O, Adea K, Hulo N, Sattonnet P, Genecand C, Iten A, et al. Infectious viral load in unvaccinated and vaccinated individuals infected with ancestral, Delta or Omicron SARS-CoV-2. Nat Med [Internet]. 2022 Apr 8 [cited 2022 Apr 20]; Available from: https://www.nature.com/articles/s41591-022-01816-0

18. Doremalen N van, Schulz JE, Adney DR, Saturday TA, Fischer RJ, Yinda CK, et al. Efficacy of ChAdOx1 vaccines against SARS-CoV-2 Variants of Concern Beta, Delta and Omicron in the Syrian hamster model [Internet]. In Review; 2022 Feb [cited 2022 Jun 9]. Available from: https://www.researchsquare.com/article/rs-1343927/v1

19. Altarawneh HN, Chemaitelly H, Hasan MR, Ayoub HH, Qassim S, AlMukdad S, et al. Protection against the Omicron Variant from Previous SARS-CoV-2 Infection. N Engl J Med. 2022 Mar 31;386(13):1288–90.

20. Carreño JM, Alshammary H, Tcheou J, Singh G, Raskin AJ, Kawabata H, et al. Activity of convalescent and vaccine serum against SARS-CoV-2 Omicron. Nature. 2022 Feb 24;602(7898):682–8.

21. Servellita V, Syed AM, Morris MK, Brazer N, Saldhi P, Garcia-Knight M, et al. Neutralizing immunity in vaccine breakthrough infections from the SARS-CoV-2 Omicron and Delta variants. Cell. 2022 Apr;185(9):1539–1548.e5.

22. Rössler A, Riepler L, Bante D, von Laer D, Kimpel J. SARS-CoV-2 Omicron Variant Neutralization in Serum from Vaccinated and Convalescent Persons. N Engl J Med. 2022 Feb 17;386(7):698–700.

23. Lyngse FP, Mortensen LH, Denwood MJ, Christiansen LE, Møller CH, Skov RL, et al. SARS-CoV-2 Omicron VOC Transmission in Danish Households [Internet]. Infectious Diseases (except HIV/AIDS); 2021 Dec [cited 2022 Jul 11]. Available from: http://medrxiv.org/lookup/doi/10.1101/2021.12.27.21268278

24. Munster VJ, Feldmann F, Williamson BN, van Doremalen N, Pérez-Pérez L, Schulz J, et al. Respiratory disease in rhesus macaques inoculated with SARS-CoV-2. Nature. 2020/05/13 ed. 2020 Sep;585(7824):268–72.

25. van Doremalen N, Purushotham JN, Schulz JE, Holbrook MG, Bushmaker T, Carmody A, et al. Intranasal ChAdOx1 nCoV-19/AZD1222 vaccination reduces shedding of SARS-CoV-2 D614G in rhesus macaques [Internet]. Microbiology; 2021 Jan [cited 2021 Mar 1]. Available from: http://biorxiv.org/lookup/doi/10.1101/2021.01.09.426058

26. Haddock E, Feldmann F, Shupert WL, Feldmann H. Inactivation of SARS-CoV-2 Laboratory Specimens. Am J Trop Med Hyg. 2021 Apr 20;104(6):2195–8.

27. Fukushi S, Mizutani T, Saijo M, Matsuyama S, Miyajima N, Taguchi F, et al. Vesicular stomatitis virus pseudotyped with severe acute respiratory syndrome coronavirus spike protein. Journal of General Virology. 2005 Aug 1;86(8):2269–74.

28. Letko M, Marzi A, Munster V. Functional assessment of cell entry and receptor usage for SARS-CoV-2 and other lineage B betacoronaviruses. Nat Microbiol. 2020 Apr;5(4):562–9.

29. Corman VM, Landt O, Kaiser M, Molenkamp R, Meijer A, Chu DK, et al. Detection of 2019 novel coronavirus (2019-nCoV) by real-time RT-PCR. Eurosurveillance [Internet]. 2020 Jan 23 [cited 2020 Dec 4];25(3). Available from: https://www.eurosurveillance.org/content/10.2807/1560-7917.ES.2020.25.3.2000045

30. Rothe C, Schunk M, Sothmann P, Bretzel G, Froeschl G, Wallrauch C, et al. Transmission of 2019-nCoV Infection from an Asymptomatic Contact in Germany. The New England journal of medicine. 2020;

